# mRNA-1273 and Ad26.COV2.S vaccines protect against the B.1.621 variant of SARS-CoV-2

**DOI:** 10.1101/2021.12.29.474432

**Authors:** Tamarand L. Darling, Baoling Ying, Bradley Whitener, Laura A. VanBlargan, Traci L. Bricker, Chieh-Yu Liang, Astha Joshi, Gayan Bamunuarachchi, Kuljeet Seehra, Aaron J. Schmitz, Peter J. Halfmann, Yoshihiro Kawoaka, Sayda M. Elbashir, Darin K. Edwards, Larissa B. Thackray, Michael S. Diamond, Adrianus C. M. Boon

**Affiliations:** Department of Medicine, Washington University School of Medicine in St. Louis, MO 63110, USA; Department of Pathology and Immunology, Washington University School of Medicine in St. Louis, MO 63110, USA; Department of Microbiology, Washington University School of Medicine in St. Louis, MO 63110, USA; The Andrew M. and Jane M. Bursky Center for Human Immunology and Immunotherapy Programs, Washington University School of Medicine. St. Louis, MO 63110, USA; Department of Pathobiological Sciences, School of Veterinary Medicine, University of Wisconsin, Madison, WI 53711; Department of Virology, Institute of Medical Science, University of Tokyo, Tokyo 108-8639, Japan; The Research Center for Global Viral Diseases, National Center for Global Health and Medicine Research Institute, Tokyo 162-8655, Japan; Moderna, Inc., Cambridge, MA 02139, USA

## Abstract

Since the emergence of severe acute respiratory syndrome coronavirus 2 (SARS-CoV-2) in 2019, viral variants with greater transmissibility or immune evasion properties have arisen, which could jeopardize recently deployed vaccine and antibody-based countermeasures. Here, we evaluated in mice and hamsters the efficacy of preclinical non-GMP Moderna mRNA vaccine (mRNA-1273) and the Johnson & Johnson recombinant adenoviral-vectored vaccine (Ad26.COV2.S) against the B.1.621 (Mu) South American variant of SARS-CoV-2, which contains spike mutations T95I, Y144S, Y145N, R346K, E484K, N501Y, D614G, P681H, and D950N. Immunization of 129S2 and K18-human ACE2 transgenic mice with mRNA-1273 vaccine protected against weight loss, lung infection, and lung pathology after challenge with B.1.621 or WA1/2020 N501Y/D614G SARS-CoV-2 strain. Similarly, immunization of 129S2 mice and Syrian hamsters with a high dose of Ad26.COV2.S reduced lung infection after B.1.621 virus challenge. Thus, immunity induced by mRNA-1273 or Ad26.COV2.S vaccines can protect against the B.1.621 variant of SARS-CoV-2 in multiple animal models.

## INTRODUCTION

Severe acute respiratory syndrome coronavirus 2 (SARS-CoV-2), the etiological agent of coronavirus disease 2019 (COVID-19), has caused hundreds of millions of infections worldwide with more than 5 million deaths. Vaccines targeting the SARS-CoV-2 spike protein were developed within one year of the start of the pandemic. Several of these (mRNA and adenoviral-vectored) are remarkably effective in protecting against severe COVID-19, with efficacy rates ranging from 75 to 95% depending on the vaccine and age of the individual [1-3]. Vaccines also protect against infection, and likely transmission, albeit at lower 50-70% rates [4, 5]. The emergence of several SARS-CoV-2 variants with amino acid substitutions in the spike protein has jeopardized the efficacy of current vaccines to protect against infection and disease. These variants can be more transmissible and also evade serum neutralizing antibodies. As an example, while the Delta variant (B.1.617.2) had no appreciable effect on vaccine efficacy against hospitalization, vaccine-mediated protection against infection was markedly reduced [1-5]. Thus, evaluating vaccine efficacy against emerging variants of SARS-CoV-2 is important for deciding when to administer booster shots, and determining if and when mono-or multivalent vaccines with variant spike antigens are needed.

After the emergence of the first D614G variant, several variants of concern (VOC) or interest (VOI) arose including B.1.1.7 (Alpha), B.1.351 (Beta), B.1.1.28 (Gamma), B.1.617.2 (Delta), and more recently B.1.1.529 (Omicron). Isolates from these lineages showed increased resistance to neutralizing antibodies and enhanced transmissibility compared to the antecedent SARS-CoV-2 strains [6-8]. Beyond these major VOC, other variants have emerged. The B.1.621 (Mu) variant was first detected in Colombia in January of 2021, and since then has spread to 51 countries including the United States, Japan, and the United Kingdom. The spike protein of B.1.621 varies at nine positions compared to the original SARS-CoV-2 isolate: T38I, Y144T, Y145S, R346K, E484K, N501Y, D614G, P681H, D950N [9]. The E484K mutation, also found in the B.1.351 (Beta) and P.1 (Gamma) variant, is predicted to reduce serum neutralizing antibody titers against this virus. The R346K mutation, first identified in B.1.621, and more recently in a subset of B.1.1.529 (Omicron), is considered a key mutation that confers resistance to serum antibodies from convalescent and vaccinated individuals [10] and class 2 neutralizing monoclonal antibodies (mAbs) [11]. In serum from individuals immunized with Ad26.COV2.S, mRNA-1273 or BNT162b2 vaccines, difference in neutralization comparing historical and B.1.621 SARS-CoV-2 ranged from 2.1-8.7 fold depending on the study population and vaccine [12-16]. A similar difference (4.7-12 fold) in neutralization was observed in convalescent sera from previously infected individuals [13, 14]. However, the impact of the spike mutations in B.1.621 on the protective efficacy of vaccines *in vivo* remains unknown. Here, we evaluated the immunogenicity and efficacy of two vaccines currently under Emergency Use Authorization (EUA), mRNA-1273 (Moderna) and Ad26.COV2.S (Johnson & Johnson), against the B.1.621 variant of SARS-CoV-2. We show that immunity induced by mRNA-1273 and Ad26.COV2.S protects mice and hamsters from challenge with the B.1.621 variant of SARS-CoV-2.

## RESULTS

### Immunogenicity and protection by Ad26.COV2-S vaccine against B.1.621 challenge in 129S2 mice

Groups of 5-6 week-old male 129S2 mice were immunized once with 10^8^, 10^9^ or 10^10^ virus particles of the Ad26.COV2.S vaccine (**Fig 1A**). Serum was collected 21 and 115 days later, and antibody responses were evaluated by ELISA for spike (S)-specific antibody responses. As expected, serum from control mice that received PBS did not bind to the S protein by ELISA (**Fig 1B**). In comparison, serum collected from mice 115 days after immunization with 10^8^, 10^9^ or 10^10^ dose of Ad26.COV2.S contained anti-S specific antibodies with a geometric mean titer (GMT) of 1:21,087, 1:19,870 and 1:80,599 respectively (**Fig 1B**). The serum anti-S antibody response was higher in animals that received 10^10^ dose (*P* < 0.001, **Fig 1B**) than those immunized with 10^8^ or 10^9^ dose of vaccine. A comparison of the anti-S response between 21 and 115 days after immunization revealed an approximately 3-fold increase in anti-S response over time (**Fig S1C**). Serum samples also were tested for neutralization of SARS-CoV-2 by focus reduction neutralization test (FRNT). Whereas serum from the control animals did not neutralize WA1/2020 N501Y/D614G or B.1.621 (**Fig 1C**), serum from mice immunized with 10^8^, 10^9^, or 10^10^ dose of Ad26.COV2.S did (WA1/2020 N501Y/D614G: GMT of 1:3,602, 1:6,071 and 1:20,592, respectively; and B.1.621: GMT of 1:3,172, 1:4,460 and 1:20,520, respectively) (**Fig 1C** and **S2A**). No significant difference (*P* > 0.5) in serum neutralization was observed between the WA1/2020 N501Y/D614G and B.1.621 strains. Also, no significant difference (*P* > 0.5) was observed in a pairwise comparison of serum neutralization titers against WA1/2020 N501Y/D614G and B.1.621 (**Fig S2A**).

**Figure 1.**
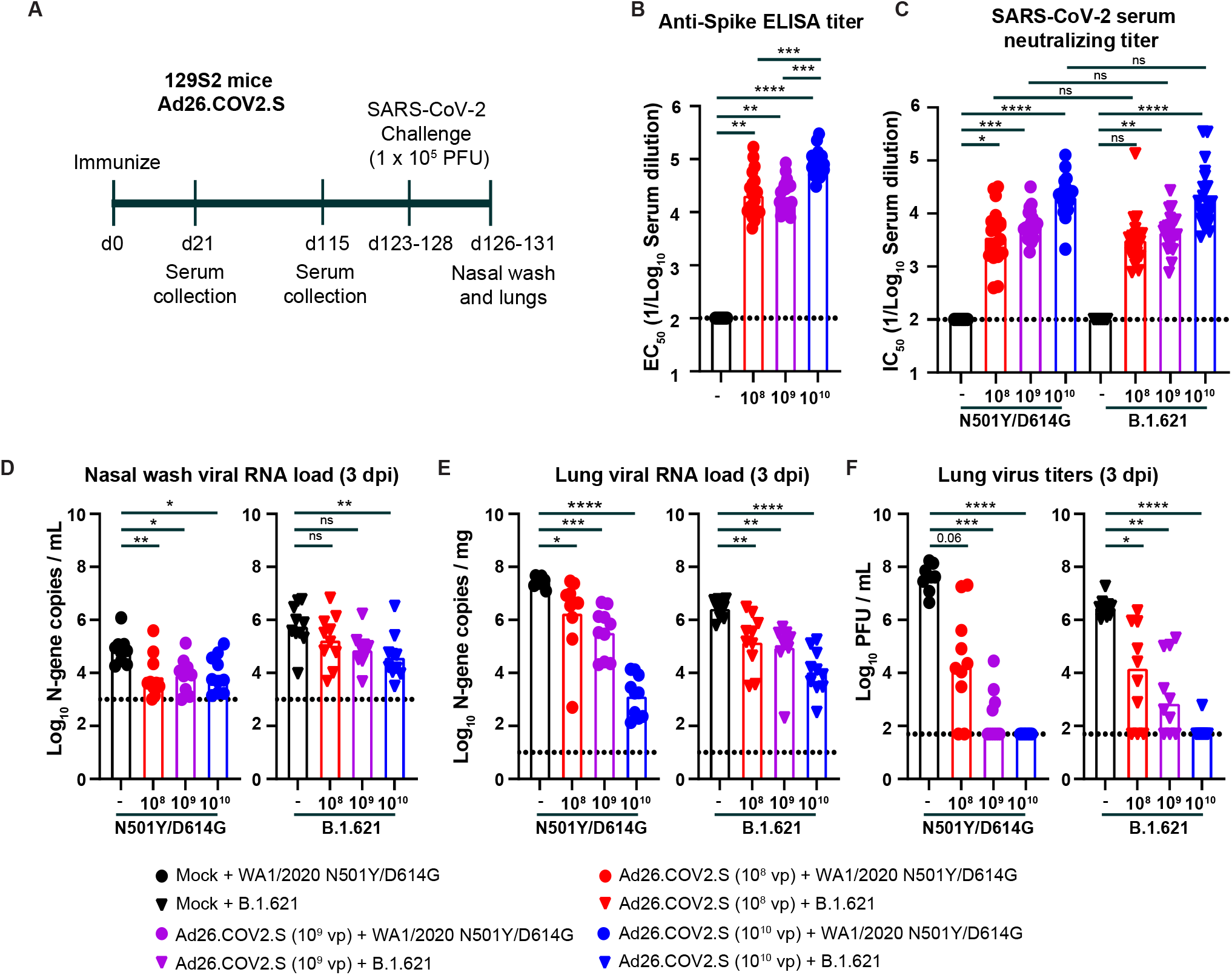
Ad26.COV2.S vaccine protects 129S2 mice against challenge with WA1/2020 N501Y/D614G and B.1.621. (**A**) Experimental setup. (**B**) Serum anti-S protein antibody response (EC_50_) in control mice (black symbols), and mice immunized with 10^8^ (red symbols), 10^9^ (purple symbols), or 10^10^ (blue symbols) of Ad26.COV2.S (**** *P* < 0.0001, *** *P* < 0.001 by non-parametric one-way ANOVA with a Dunn’s post-test). (**C**) Serum neutralizing titer (IC_50_) against WA1/2020 N501Y/D614G (circles) or B.1.621 (triangles) measured by FRNT from 129S2 mice immunized once with 10^8^ (red symbols), 10^9^ (purple symbols), or 10^10^ (blue symbols) of Ad26.COV2.S (**** *P* < 0.0001, *** *P* < 0.001, ns = not significant by non-parametric one-way ANOVA with a Dunn’s post-test). (**D-F**) 129S2 mice were challenged with 10^5^ PFU of the WA1/2020 N501Y/D614G (circles) or B.1.621 (triangles) variant of SARS-CoV-2, and nasal washes (**D**) and lungs (**E-F**) were collected for analysis of viral RNA levels by RT-qPCR (**D**) and infectious virus by plaque assay (**E-F**) (**** *P* < 0.0001, *** *P* < 0.001, ** *P* < 0.01, * *P* < 0.05, ns = not significant by one-way ANOVA with a Dunnett’s (**D-E**) or Dunn’s (**F**) post-test). (**B-G**) Bars indicate the geometric mean values, and dotted lines are the LOD of the assays. The results are from two independent experiments, and each symbol represents an individual animals.

Next, we challenged the Ad26.COV2.S-immunized 129S2 mice with 10^5^ plaque forming-units (PFU) of WA1/2020 N501Y/D614G or B.1.621 virus, and 3 days later collected nasal washes and the left lung lobe for viral burden analysis. We used 129S2 mice for these studies because these animals are permissive to infection by some SARS-CoV-2 variants (*e*.*g*., B.1.1.7, B.1.1.28, and B.1.351) or mouse-adapted or engineered strains [17-19] that encode an N501Y mutation, which enables engagement of murine ACE2 [20]. Infection of 129S2 mice with SARS-CoV-2 results in mild to moderate lung infection and clinical disease with subsequent recovery [17, 19]. In the nasal wash of control animals challenged with WA1/2020 N501Y/D614G, we detected ∼10^5^ copies of the *N* gene transcript per mL (**Fig 1D**). Immunization with 10^8^, 10^9^, or 10^10^ dose of Ad26.COV2.S reduced the viral RNA levels by 10, 9 and 8-fold (*P* < 0.01, 0.05, and 0.05 respectively, **Fig 1D**). After challenge with B.1.621, we measured ∼10^6^ copies of *N* gene transcript per mL in the nasal wash, and this was reduced 3, 8, and 15-fold (*P* < 0.01 for 10^10^ dose) in animals immunized with 10^8^, 10^9^, or 10^10^ dose of Ad26.COV2.S, respectively.

Viral RNA levels also were quantified in the left lung lobe at 3 dpi. In the control groups challenged with the WA1/2020 N501Y/D614G virus, ∼10^7^ *N* gene transcript copies per mg and ∼10^7^ PFU/mL were measured in lung homogenates (**Fig 1E-F**). Immunization with the 10^8^, 10^9^, or 10^10^ dose of Ad26.COV2.S reduced the *N* gene copy number by 15, 80, and 20,000-fold (*P* < 0.05, 0.001, and 0.0001 respectively) and infectious virus levels by 10^3^, 10^5^, and 10^6^-fold (*P* = 0.06, < 0.001, and < 0.0001, respectively) (**Fig 1E-F**). A second cohort of Ad26.COV2.S-immunized animals were challenged with the B.1.621 strain of SARS-CoV-2. In the control groups, we detected ∼10^6^ *N* gene transcript copies per mg and ∼10^6^ PFU/mL of infectious virus of lung homogenate (**Fig 1E-F**). Immunization with the 10^8^, 10^9^, or 10^10^ dose of Ad26.COV2.S reduced the viral RNA levels by 18, 30, and 250-fold (*P* < 0.01, 0.01, and 0.0001, respectively) and infectious virus burden by 350, 2,000, and 50,000-fold (*P* < 0.05, 0.01, and 0.0001, respectively) (**Fig 1E-F**).

### Immunogenicity and protection by mRNA-1273 against B.1.621 challenge in K18-hACE2 mice

Next, we evaluated the efficacy of preclinical non-GMP lots of the Moderna mRNA-1273 vaccine encoding a sequenced-optimized 2 proline-stabilized spike protein of Wuhan-1 for protection against B.1.621 in K18-hACE2 transgenic mice. Groups of 7-8 week-old female mice were immunized and boosted via intramuscular route with a low (0.25 μg) or high (5 μg) dose of the mRNA-1273 or a control mRNA vaccine (mRNA-control, **Fig 2A**). Serum was collected 21 days after the second immunization, and inhibitory antibody responses were evaluated by FRNT against WA1/2020 N501Y/D614G and B.1.621. As expected, serum from mice immunized with the control vaccine did not inhibit virus infection (**Fig 2B**). In contrast, serum from mice immunized with 0.25 μg dose of mRNA-1273 neutralized infectious virus with GMT of 1:1,125 and 1:434 for WA1/2020 N501Y/D614G and B.1.621 viruses, respectively. Serum from mice immunized with 5 μg of mRNA-1273 showed greater neutralizing activity against WA1/2020 N501Y/D614G and B.1.621 with GMT of 1:19,751 and 1:15,130 respectively. A statistical difference in serum GMT was observed between WA1/2020 N501Y/D614G and B.1.621 for the low dose (*P* < 0.05), but not the high dose (*P* > 0.5) vaccine. However, a pairwise comparison showed a 1.3 to 2.3-fold difference in serum neutralizing antibody titer between WA1/2020 N501Y/D614G and B.1.621 for the high (*P* < 0.05) and low (*P* < 0.01) dose mRNA-1273 immunized mice (**Fig S2B**).

**Figure 2.**
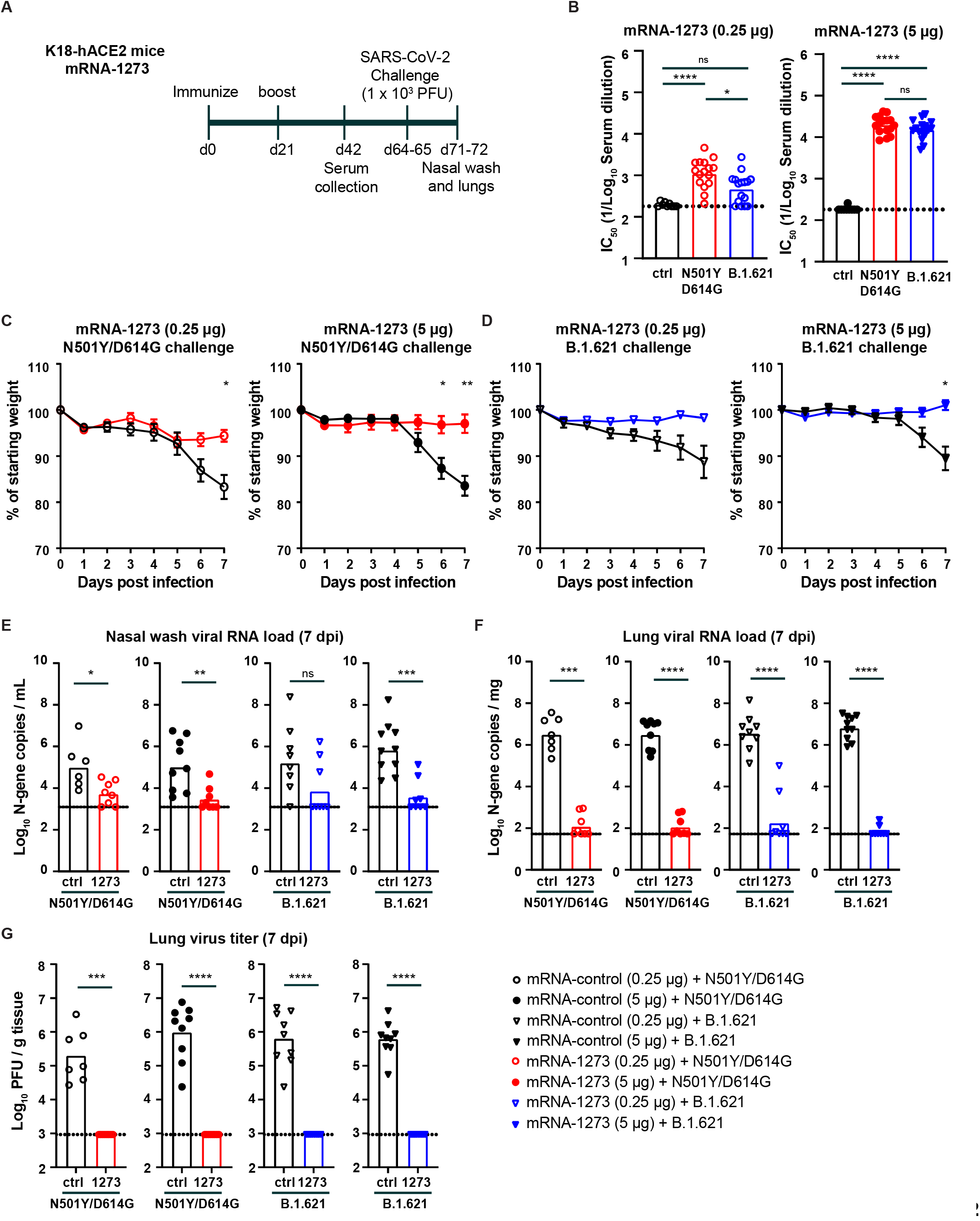
mRNA-1273 protects K18-hACE2 transgenic mice against challenge with N501Y/D614G and B.1.621. (**A**) Experimental setup. (**B**) Serum neutralizing titer (IC_50_) against WA1/2020 N501Y/D614G (red circles) or B.1.621 (blue triangles) from K18-hACE2 mice immunized twice with 0.25 μg (open symbols) or 5 μg (closed symbols) of mRNA-1273 or mRNA control vaccine (*** *P* < 0.001, **** *P* < 0.0001 by non-parametric one-way ANOVA with a Dunn’s post-test). (**C-D**) Mean ± SEM of weight loss/gain in SARS-CoV-2 challenged mice (** *P* < 0.01, * *P* < 0.05, ns = not significant by two-way ANOVA). (**E-G**) K18-hACE2 mice were challenged with 10^3^ PFU of the WA1/2020 N501Y/D614G (red circles) or B.1.621 (blue triangles), and nasal washes (**E**) and lungs (**F-G**) were collected for analysis of viral RNA levels by RT-qPCR (**E**) and infectious virus by plaque assay (**F-G**) (**** *P* < 0.0001, *** *P* < 0.001, ** *P* < 0.01, * *P* < 0.05, ns = not significant by Mann-Whitney test). (**B, E-G**) Bars indicate the geometric mean values, and dotted lines are the LOD of the assays. The results are from two independent experiments, and each symbol represents an individual animals.

We next evaluated the protective effect of mRNA-1273 vaccine in K18-hACE2 mice. Mice immunized with mRNA-1273 or control mRNA vaccine were challenged via intranasal route with 10^3^ PFU of WA1/2020 N501Y/D614G or B.1.621 virus, and body weight was recorded for 7 days before a nasal wash and the left lung lobe were collected for viral burden analysis. In the control groups challenged with the WA1/2020 N501Y/D614G virus, substantial weight loss was observed at 6 and 7 dpi. Immunization with 0.25 or 5 μg of mRNA-1273 significantly prevented weight loss (*P* < 0.05 and 0.01 respectively, **Fig 2C**). Similar protection against weight loss after challenge with B.1.621 was observed in the groups immunized with 0.25 (*P* = 0.19) or 5 μg (*P* < 0.05) of mRNA-1273 (**Fig 2D**).

We quantified the amount of viral RNA in the nasal wash at 7 dpi. In control vaccinated animals challenged with WA1/2020 N501Y/D614G, we detected ∼10^5^ copies of the SARS-CoV-2 *N* gene transcript per mL in the nasal wash (**Fig 2E**). Immunization with mRNA-1273 reduced the levels of WA1/2020 N501Y/D614G RNA by 20 to 45-fold for the 0.25 μg (*P* < 0.05) and 5 μg (*P* < 0.01) doses, respectively (**Fig 2E**). In the groups vaccinated with the control mRNA and challenged with the B.1.621 virus, we detected 10^5^-10^6^ copies of the *N* gene transcript per mL in the nasal wash (**Fig 2E**). Immunization with 0.25 μg (*P* = 0.1) or 5 μg (*P* < 0.001) of mRNA-1273 reduced the *N* gene copy number by ∼25 and 200-fold, respectively (**Fig 2E**).

The amount of virus in lung homogenates also was quantified. In the control group challenged with WA1/2020 N501Y/D614G, ∼10^6^ copies of the *N* transcript per mg and 10^5^-10^6^ PFU/g tissue were detected (**Fig 2F-G**). Immunization with 0.25 μg of mRNA-1273 resulted in markedly reduced (∼20,000-fold, *P* < 0.0001) levels of viral RNA and no detectable infectious virus titer in the lung (**Fig 2F-G**). Similar results were seen in mice immunized with 5 μg of mRNA-1273 (**Fig 2F-G**). A second group of immunized animals were challenged with B.1.621. In the control mRNA vaccinated groups, we detected high levels of viral RNA (*N* gene, >10^6^ copies per mg) and infectious virus (6 × 10^5^ PFU/g) in lung homogenates (**Fig 2F-G**). Immunization with either dose of mRNA-1273 vaccine significantly reduced levels of B.1.621 viral RNA and infectious virus in the lung (∼20,000-fold, *P* < 0.0001, **Fig 2F-G**).

K18-hACE2 mice vaccinated with 0.25 μg or 5 μg of mRNA-1273 also had markedly reduced, if not absent, lung pathology at 7 dpi compared to the control mRNA vaccinated and challenged animals (**Fig S3**). Overall, these data indicate that the mRNA-1273 vaccine protects against the B.1.621 variant of SARS-CoV-2 in K18-hACE2 mice.

### Immunogenicity and protection by mRNA-1273 vaccine against B.1.621 challenge in 129S2 mice

To corroborate our findings, we also tested the mRNA-1273 vaccine in immunocompetent 129S2 mice. Groups of 7-8 week-old female mice were immunized and boosted via intramuscular route with 0.25 or 5 μg of mRNA-1273 or control mRNA vaccine (**Fig 3A**). Serum was collected 21 days after the second immunization, and antibody responses were evaluated by FRNT. As expected, serum from 129S2 mice immunized with the control mRNA vaccine did not neutralize virus infection (**Fig 3B**). In contrast, serum from mice immunized with 5 μg of mRNA-1273 robustly neutralized infection with GMTs of 1:40,066 and 1:38,675 for WA1/2020 N501Y/D614G and B.1.621, respectively. Sera from mice immunized with the 0.25 μg dose of mRNA-1273 inhibited infection of WA1/2020 N501Y/D614G and B.1.621 to a lesser extent with GMTs of 1:2,665 and 1:2,407, respectively. No significant difference (*P* > 0.5) in vaccine-induced GMTs were observed between the WA1/2020 N501Y/D614G and B.1.621 viruses (**Fig 3B**). Also no significant difference (*P* > 0.5) was observed in a pairwise comparison of the neutralization titer against WA1/2020 N501Y/D614G and B.1.621 in serum from 129S2 mice immunized with high and low dose mRNA-1273 vaccine (**Fig S2C**).

**Figure 3.**
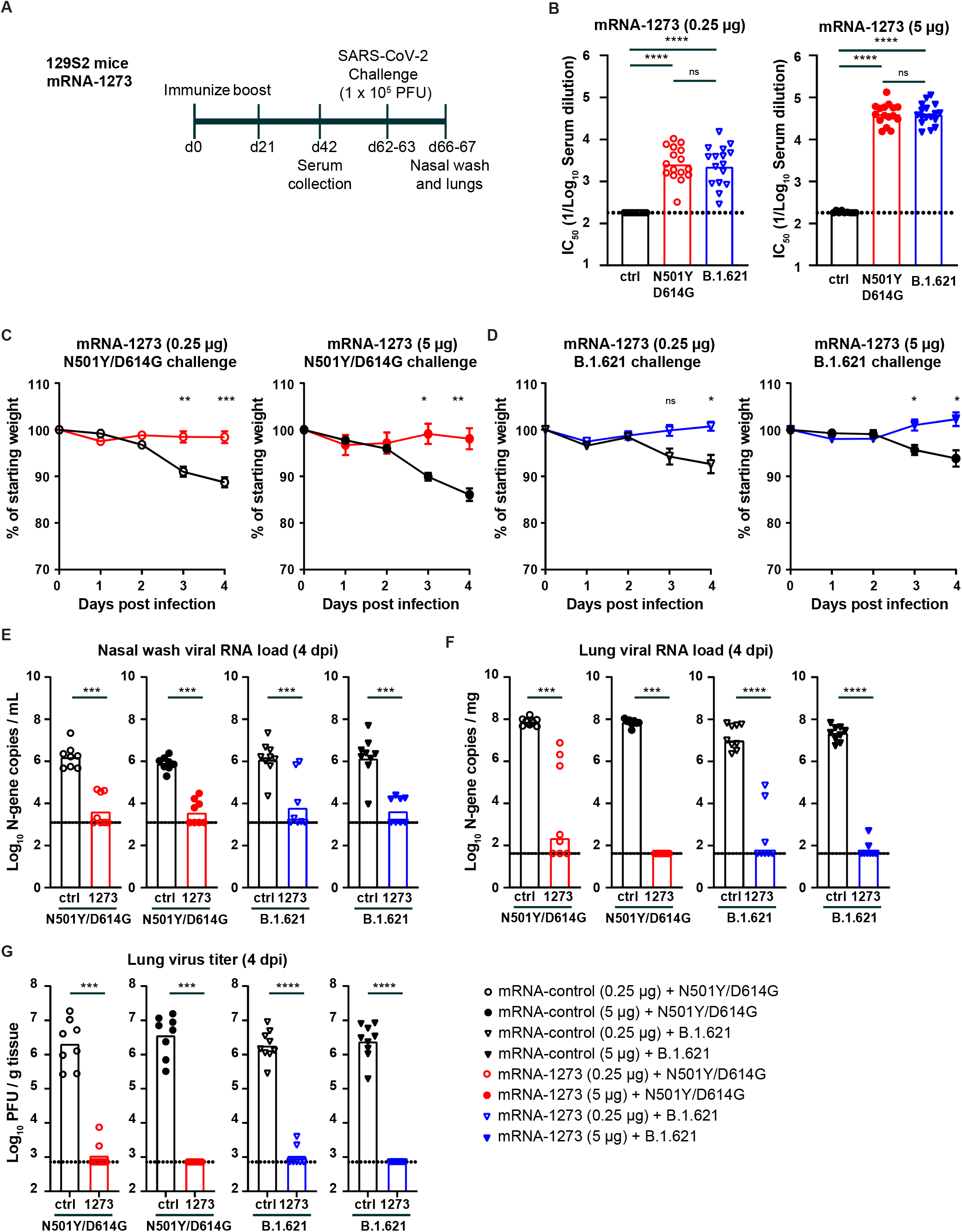
mRNA-1273 protects 129S2 mice against challenge with WA1/2020 N501Y/D614G and B.1.621. (**A**) Experimental setup. (**B**) Serum neutralizing titer (IC_50_) against WA1/2020 N501Y/D614G (red circles) or B.1.621 (blue triangles) from 129S2 mice immunized twice with 0.25 μg (open symbols) or 5 μg (closed symbols) of mRNA-1273 or mRNA control vaccine. (**** *P* < 0.0001, *** *P* < 0.001 by non-parametric one-way ANOVA with a Dunn’s post-test). (**C-D**) Mean ± SEM of weight loss/gain in SARS-CoV-2 challenged mice (** *P* < 0.01, * *P* < 0.05, ns = not significant by two-way ANOVA). (**E-G**) 129S2 mice were challenged with 10^5^ PFU of WA1/2020 N501Y/D614G (red symbols) or B.1.621 (blue symbols), and nasal washes (**E**) and lungs (**F-G**) were evaluated for viral RNA levels by RT-qPCR (**E**) and infectious virus by plaque assay (**F-G**) (**** *P* < 0.0001, *** *P* < 0.001, ** *P* < 0.01, * *P* < 0.05, ns = not significant by Mann-Whitney test). (**B, E-G**) Bars indicate the geometric mean values, and dotted lines are the LOD of the assays. The results are from two independent experiments, and each symbol represents an individual animals.

We next challenged immunized 129S2 mice with 10^5^ PFU of WA1/2020 N501Y/D614G or B.1.621 via an intranasal route. Weights were recorded for 4 days before the animals were sacrificed, and nasal washes and the left lung lobe were collected for viral burden analysis. Control animals challenged with the WA1/2020 N501Y/D614G virus lost ∼10% of their starting body weight by 4 dpi (**Fig 3C**). Immunization with 0.25 and 5 μg of mRNA-1273 reduced the weight loss (*P* < 0.001 and 0.01 respectively, **Fig 3C**). Inoculation of control-vaccinated 129S2 mice with B.1.621 resulted in ∼7% weight loss at 4 dpi (**Fig 3D**), and immunization with 0.25 or 5 μg of mRNA-1273 prevented this weight loss (*P* < 0.05).

In animals vaccinated with the control mRNA and challenged with the WA1/2020 N501Y/D614G virus, we detected ∼10^6^ copies/mL of *N* gene transcript (**Fig 3E**) in the nasal wash at 4 dpi. Immunization with 0.25 or 5 μg of mRNA-1273 reduced WA1/2020 N501Y/D614G viral RNA levels in the nasal wash (∼200-fold, *P* < 0.001). In mice vaccinated with the control mRNA vaccine and challenged with B.1.621, we detected ∼10^6^ *N* gene transcript copies per mL in the nasal wash (**Fig 3E**). Immunization with 0.25 μg or 5 μg of mRNA-1273 reduced the viral RNA levels substantially (∼200-fold, *P* < 0.001; **Fig 3E**). Viral burden also was measured in lung homogenates of these same animals. In the control mRNA group challenged with the WA1/2020 N501Y/D614G virus, we detected ∼10^8^ copies/mg of *N* gene transcript (**Fig 3F**) and > 10^6^ PFU/g of infectious virus (**Fig 3G**) in lung homogenates. Immunization with 0.25 μg of mRNA-1273 effectively reduced the viral RNA (10^6^-fold, *P* < 0.001, **Fig 3F**) and infectious virus levels (1,000-fold, *P* < 0.001, **Fig 3G**), although some animals showed breakthrough infection. Immunization with 5 μg dose of mRNA-1273 also reduced the levels of viral RNA (10^6^-fold, *P* < 0.001) and infectious virus (1,000-fold, *P* < 0.001, **Fig 3F-G**). Importantly, no breakthrough infection was detected. A second group of immunized animals were challenged with the B.1.621 virus. In the control mRNA vaccinated groups, we detected in the lung homogenates ∼10^7^ copies/mg of viral RNA transcript (**Fig 3F**) and ∼10^6^ PFU/g of infectious virus (**Fig 3E**). Immunization with 0.25 or 5 μg doses of mRNA-1273 significantly reduced B.1.621 viral RNA (10^5^-fold, *P* < 0.0001) and infectious virus (1,000-fold, *P* < 0.0001) levels in the lung (**Fig 3F-G**) to a similar degree after WA1/2020 N501Y/D614G challenge.

The reduction in viral burden in the lungs of mice immunized with 0.25 or 5 μg of mRNA-1273 and challenged with WA1/2020 N501Y/D614G or B.1.621 corresponded with an absence of lung pathology at 4 dpi (**Fig S4**). Lung sections from mice immunized with a control mRNA and challenged with WA1/2020 N501Y/D614G or B.1.621 showed evidence of immune cell infiltration and tissue damage. Immunization with 0.25 and 5 μg of mRNA-1273 prevented the lung pathology after challenge with both WA1/2020 N501Y/D614G and B.1.621 virus. Overall, these data indicate that the mRNA-1273 vaccine protects against the B.1.621 variant of SARS-CoV-2 in non-transgenic immunocompetent 129S2 mice.

### Immunogenicity and protection by Ad26.COV2-S vaccine against B.1.621 challenge in Syrian hamsters

We next evaluated the efficacy of a single dose of the Ad26.COV2.S vaccine in Syrian hamsters. Syrian hamsters are naturally susceptible to SARS-CoV-2 and considered an excellent model for COVID-19 [19, 21-24]. Groups of 5-6 week-old male hamsters were immunized once with 10^8^ or 10^10^ dose of the Ad26.COV2.S vaccine (**Fig 4A**). Serum was collected 21 days later, and antibody responses were evaluated by ELISA and FRNT. As expected, serum from control hamsters that received PBS did not bind viral S protein (**Fig 4B**). However, sera collected from hamsters immunized with 10^8^ or 10^10^ dose of Ad26.COV2.S contained anti-S antibodies with GMTs of 1:7,506, and 1:40,913 respectively (**Fig 4B**). The serum anti-S antibody response was higher in animals receiving the 10^10^ vaccine dose (*P* < 0.05, **Fig 4B**). Serum samples were tested for neutralization of SARS-CoV-2 by FRNT. Serum from hamsters immunized with 10^8^ or 10^10^ dose of Ad26.COV2.S neutralized WA1/2020 with GMT of 1:381 and 1:1,692 respectively (**Fig 4C**), and B.1.621 with GMT of 1:136 and 1:501, respectively (**Fig 4C**). No difference in GMT was observed with respect to neutralization of WA1/2020 and B.1.621 variant after immunization with 10^8^ *(P* = 0.06) or 10^10^ (*P* = 0.6) dose of Ad26.COV2.S (**Fig 4C**). A pair-wise comparison found a ∼3-fold reduction in serum neutralizing antibody titer between WA1/2020 and B.1.621 for the high (*P* < 0.0001) and low (*P* < 0.001) dose Ad26-COV2.S immunized hamsters (**Fig S2D**).

**Figure 4.**
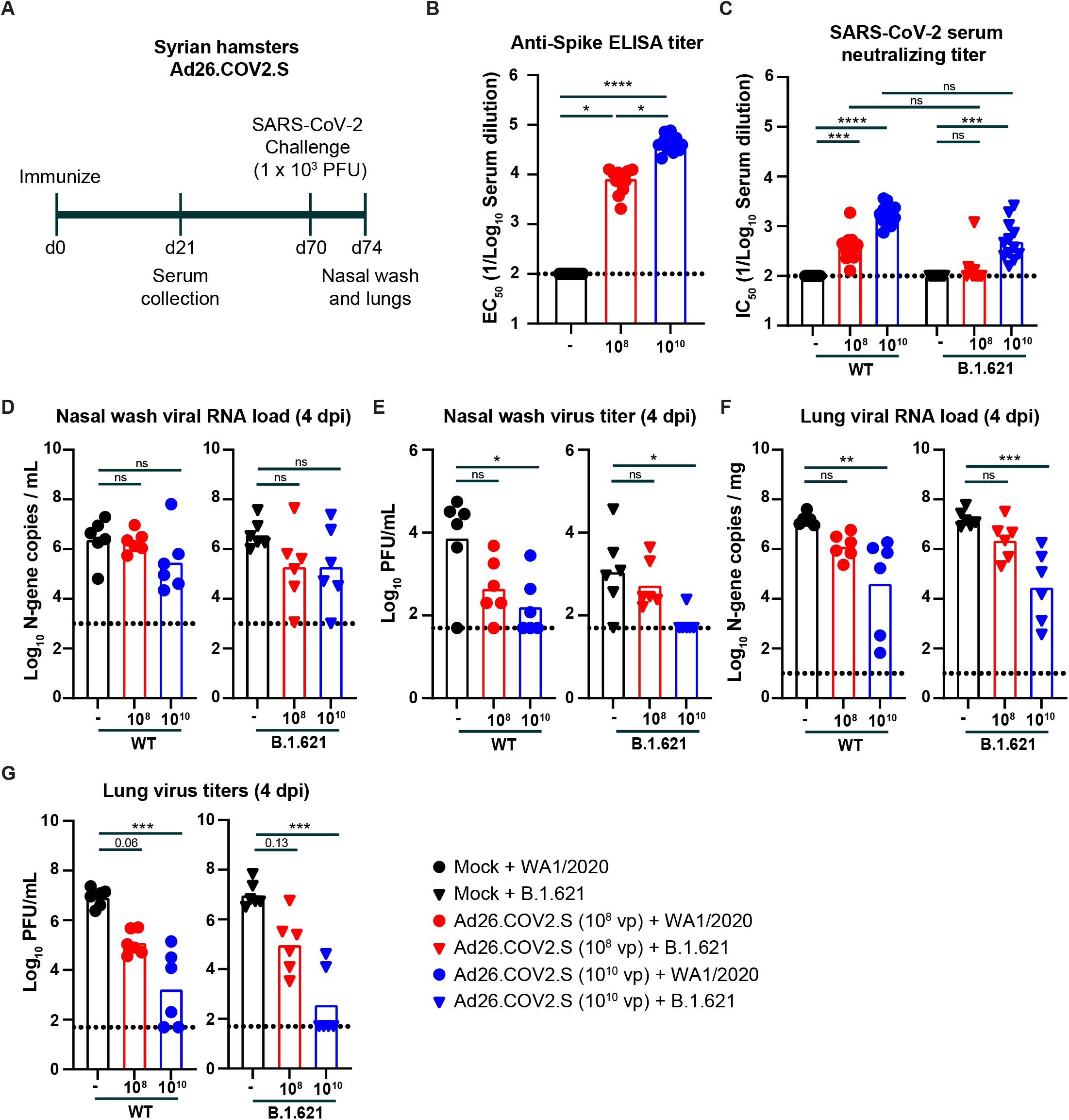
Ad26.COV2.S vaccine protects Syrian hamsters against challenge with WA1/2020 and B.1.621. (**A**) Experimental setup. (**B**) Serum anti-S protein antibody response (EC_50_) in control hamsters (black symbols), and hamsters immunized with 10^8^ (red symbols) or 10^10^ (blue symbols) of Ad26.COV2.S (**** *P* < 0.0001, *** *P* < 0.001, ns = not significant by non-parametric one-way ANOVA with a Dunn’s post-test. (**C**) Serum neutralizing titer (IC_50_) against WA1/2020 (circles) or B.1.621 (triangles) from hamsters immunized once with 10^8^ (red symbols) or 10^10^ (blue symbols) of Ad26.COV2.S (**** *P* < 0.0001, *** *P* < 0.001, ns = not significant by non-parametric one-way ANOVA with a Dunn’s post-test. (**D-G**) Syrian hamsters were challenged with 10^3^ PFU of the WA1/2020 (circles) or B.1.621 (triangles), and nasal washes (**D-E**) and lungs (**F-G**) were evaluated for viral RNA levels by RT-qPCR (**D** and **F**) and infectious virus by plaque assay (**E** and **G**) (**** *P* < 0.0001, *** *P* < 0.001, ** *P* < 0.01, * *P* < 0.05, ns = not significant by one-way ANOVA with a Dunnett’s (**D** and **F**) or Dunn’s (**E** and **G**) post-test). (**B-G**) Bars indicate the geometric mean values, and dotted lines are the LOD of the assays. The results are from one experiment, and each symbol represents an individual animal.

Seventy days after immunization, the hamsters were challenged via an intranasal route with 10^3^ PFU of WA1/2020 or B.1.621. Four days later, the animals were sacrificed, and a nasal wash and the left lung lobe were collected for viral burden analysis. In the nasal wash of control animals challenged with the WA1/2020 virus, we detected ∼10^6^ copies of the *N* gene transcript per mL of nasal wash (**Fig 4D**) and ∼10^4^ PFU/mL of infectious virus (**Fig 4E**). Immunization with 10^8^ or 10^10^ dose of Ad26.COV2.S did not reduce the viral RNA levels in the nasal wash (**Fig 4D**), although the infectious virus titer was decreased by 16 and 46-fold (*P* > 0.05 and < 0.05 respectively, **Fig 4E**). Upon challenge of hamsters with B.1.621, we detected ∼10^6^ *N* gene copies per mL (**Fig 4D**) and ∼10^3^ PFU/mL of infectious virus (**Fig 4E**) in the nasal wash. Immunization with 10^8^ or 10^10^ dose of Ad26.COV.2 reduced the *N* gene copy number 19-fold (*P* > 0.05, **Fig 4D**). In comparison, the infectious virus titer was reduced by 2 and 17-fold (*P* > 0.05 and < 0.05 for 10^8^ and 10^10^ dose respectively, **Fig 4E**)

We also measured the viral burden in lung homogenates of hamsters. In the control group challenged with WA1/2020, the *N* gene copy number was ∼10^7^ copies per mg (**Fig 4F**) and the infectious virus levels were ∼10^7^ PFU/mL (**Fig 4G**). Immunization with the 10^8^ or 10^10^ dose of Ad26.COV2.S reduced the viral RNA (11 and 360-fold, *P* > 0.05 and < 0.01) and infectious virus (66 and 4800-fold, *P* = 0.06 and < 0.001) levels (**Fig 4F-G**). After B.1.621 challenge of the control group, we detected ∼10^7^ *N* gene copies per mg of tissue (**Fig 4F**) and ∼10^7^ PFU/mL of infectious virus in the lung (**Fig 4G**). Immunization with the 10^8^ or 10^10^ dose of Ad26.COV2.S reduced the viral RNA (7 and 540-fold, *P* > 0.05 and < 0.05) and infectious virus (100 and 25,500-fold, *P* = 0.13 and < 0.001) levels (**Fig 4F-G**).

## DISCUSSION

In this study, we evaluated the efficacy of two vaccines under EUA, mRNA-1273 and Ad26.COV2.S, against the B.1.621 variant of SARS-CoV-2 in in three pre-clinical models; 129S2-immuno-competed mice, K18-hACE2 transgenic mice, and Syrian hamster. The mRNA-1273 vaccine induced high levels of neutralizing antibodies against WA1/2020 N501Y/D614G and B.1.621 viruses, and this response was associated with robust protection from an intranasal challenge. Immunization of 129S2 mice or Syrian hamsters with different doses of Ad26.COV2.S induced moderate to high serum neutralizing antibody responses against the B.1.621 virus. However, only the high dose (10^10^ virus particles) Ad26.COV2.S reduced virus titers substantially.

Our studies provide a comparison of the immunogenicity and efficacy of the Ad26.COV2.S and mRNA-1273 vaccines in 129S2 mice. The serum neutralizing antibody titer was similar after one dose of Ad26.COV2.S or two doses of mRNA-1273 in129S2 mice, and this was true for both the high dose (5 μg vs. 10^10^ virus particles) and the low dose (0.25 μg vs. 10^8^ virus particles) vaccine regimen. Despite the similarity in neutralization titer, the mRNA-1273 vaccine more effectively reduced viral load in the lungs of WA1/2020 N501Y/D614G or B.1.621 challenged animals than the Ad26.COV2.S vaccine. In mice immunized with a low-dose of the mRNA-1273 vaccine, approximately 20% of the animals showed evidence of breakthrough infections after challenge (**Fig 3G**). In contrast, 75% of the animals that received a low dose of Ad26.COV2.S and were challenged with WA1/2020 N501Y/D614G or B.1.621 showed virus breakthrough despite relatively equivalent levels of serum neutralizing antibodies titers at the time of challenge (**Fig 1F**). One explanation for this difference could that the mRNA vaccine requires two doses, while the Ad26.COV2.S vaccine was only given once. Another possibility is the time between the last immunization and the virus challenge, which is 41 and 115 days for the mRNA-1273 and Ad26.COV2.S immunized animals, respectively. Differences in the glycosylation pattern or the IgG subclass of antibodies between anti-S antibodies induced by mRNA-1273 and Ad26.COV2.S induced could contribute to differences in protection, as seen in non-human primates and humans [25, 26]. It is also possible that mRNA-1273 vaccine induced a better anamnestic B or T cell response in 129S2 mice compared to the Ad26.COV2.S vaccine at the time of virus challenge.

The antibody response after vaccination varied between the mouse and hamster models. Immunization of mice with 5 μg of mRNA-1273 or 10^10^ Ad26.COV2.S induced serum neutralizing antibody responses with a GMT of > 10,000 in both 129S2 and K18-hACE2 mice. In contrast, in Syrian hamsters, the GMT against WA1/2020 was ∼10-fold lower than in mice, yet still several fold higher than that observed in humans vaccinated with one dose of Ad26.COV2.S [27-30]. The reason for this difference in vaccine response between mice and hamsters remains unknown. It is possible that the hamster immune response targets different epitopes on the spike protein. Alternatively, the spike protein contains fewer T cell epitopes for hamsters compared to the mouse, although that seems unlikely given the size of the antigen. The Syrian hamster also may be a more tolerogenic, perhaps due to its complex microbiome as opposed to the SPF microbiome of 129S2 and K18 TG mice. Mice that received the microbiome from pet stores or from field mice had blunted vaccine responses compared to laboratory-housed mice in pathogen-free facilities [31]. Additional studies are required to elucidate the causes for this difference, but in general, the magnitude of antibody response in Syrian hamsters is more similar to the human antibody response after vaccination.

The B.1.621 (Mu) variant of SARS-CoV-2 has R346K and E484K mutations in the receptor binding domain of the spike protein and is believed to be more resistant to virus neutralization by serum antibodies compared to the historical SARS-CoV-2 virus. In sera from vaccinated or infected individuals, the fold difference in neutralization between the D614G (B.1) variant of SARS-CoV-2 and B.1.621 was between 2 and 12-fold [12-16, 32]. In Syrian hamsters, we observed a ∼3-fold decrease (*P* < 0.001) in serum neutralization titer between the WA1/2020 and B.1.621 virus. In K18-hACE2 mice immunized with mRNA-1273 the difference in serum neutralization titer between WA1/2020 N501Y/D614G and B.1.621 was 1.3 to 2.3-fold. In contrast, no difference in neutralization titer between WA1/2020 N501Y/D614G and B.1.621 was observed in 129S2 mice immunized with mRNA-1273 or Ad26.COV2.S, suggesting that mouse strain and species-specific differences in the antibody response. Another reason may be the N501Y mutation in WA1/2020 N501Y/D614G, which has been shown to reduce the neutralization titer of serum and certain monoclonal antibodies compared to WA1/2020 and WA1/2020 D614G, respectively [6, 33]. We used the WA1/2020 N501Y/D614G virus because the N501Y mutation was required for virus infection in immunocompetent 129S2 mice lacking hACE2 expression. Finally, it is possible that the insertion of a threonine at position 144-145 in our particular B.1.621 isolate reduced the resistance to serum neutralizing antibodies.

We observed no difference in efficacy of the mRNA and adenoviral-vectored vaccine to protect against B.1.621 in three different animal models. Near full protection, defined by undetectable levels of SARS-CoV-2 viral RNA and infectious virus, plus the absence of immunopathology in the vaccinated animals, was observed against both B.1.621 and the control WA1/2020 virus in mice immunized with mRNA-1273. While immunization with lower doses of Ad26.COV2.S offered only partial protection, we did not observe a difference in virus titer or frequency of breakthrough infection between B.1.621 and control virus. This suggests that a 2-3-fold reduction in serum neutralization titer has limited impact on mRNA and adenoviral vaccine protection against the variant B.1.621 virus.

### Limitations of the study

We note several limitations of our study. (a) We did not evaluate the effects of the vaccine on the transmission of SARS-CoV-2 in Syrian hamsters, which may be an important measure of vaccine protection. (b) We used lower doses of vaccine to mimic suboptimal and possibly waning immunity. Studies that directly compare the quality of a waning immune response to that of a low dose vaccine induced immune response are needed. (c) The challenge dose of SARS-CoV-2 used in our hamster model (10^3^ PFU) is several orders of magnitude higher that the minimal infectious dose (5 PFU) [34]. While this creates a very robust virus challenge model, it could underestimate the protective effects of vaccines. (d) We did not establish correlates of immune protection. We noted that lower serum antibody neutralization titers were associated with high viral loads and infectious virus titers in the Ad26.COV2.S immunized animals, as well as breakthrough infections in some of the mRNA-1273 vaccinated mice, albeit this did not explain all breakthrough infections. A more detailed analysis of T cell and non-neutralizing antibody responses coupled with even lower vaccine doses may be needed to fully establish a correlate of protection against breakthrough infection.

Overall, our studies demonstrate that the Moderna mRNA-1273 and Johnson & Johnson Ad26.COV2.S vaccines authorized for emergency use are immunogenic in mice and Syrian hamsters and protect against the B.1.621 (Mu) variant of SARS-CoV-2 without substantial loss of potency.

## ACKNOWLEDGEMENTS

We thank Florian Krammer and Ali Ellebedy for the plasmid and recombinant S protein respectively. This study was supported by the NIH (R01 AI157155, U01 AI151810, NIAID Centers of Excellence for Influenza Research and Response (CEIRR) contract 75N93021C00014 and 75N93021C00016, and the Collaborative Influenza Vaccine Innovation Centers (CIVIC) contract 75N93019C00051). It was also supported, in part, by the National Institutes of Allergy and Infectious Diseases Center for Research on Influenza Pathogenesis (HHSN272201400008C), and the Japan Program for Infectious Diseases Research and Infrastructure (JP21wm0125002) from the Japan Agency for Medical Research and Development (AMED).

## AUTHOR CONTRIBUTIONS

T.L.B., B.Y., B.W., C-Y.L., L.A.V., G.B., and T.L.D. performed mouse and hamster experiments. T.L.B., B.Y., and T.L.D. quantified virus titers in collected tissues. K.S., A.J., and B.W. determined viral load by real-time quantitative RT-PCR. Y.K. and P.J.H. isolated, expanded and sequenced the B.1.621 virus. T.L.D. and L.A.V. performed the virus neutralization assays. M.S.D., A.C.M.B., T.L.D. L.B.T. and B.Y. analyzed the data. A.C.M.B., and M.S.D. wrote the manuscript, and all authors edited the final version.

## DECLARATION OF INTERESTS

The Boon laboratory has received unrelated funding support in sponsored research agreements from AI Therapeutics, GreenLight Biosciences Inc., and Nano targeting & Therapy Biopharma Inc. The Boon laboratory has received funding support from AbbVie Inc., for the commercial development of SARS-CoV-2 mAb. M.S.D. is a consultant for Inbios, Vir Biotechnology, Senda Biosciences, and Carnival Corporation, and on the Scientific Advisory Boards of Moderna and Immunome. The Diamond laboratory has received unrelated funding support in sponsored research agreements from Vir Biotechnology, Kaleido, and Emergent BioSolutions and past support from Moderna not related to these studies. S.E. and D.K.E. are employees of and shareholders in Moderna Inc.

## STAR METHODS

### RESOURCE AVAILABILITY

#### Lead contact

Further information and requests for resources and reagents should be directed to the Lead Contact, Adrianus C.M. Boon.

#### Materials availability

All requests for resources and reagents should be directed to the Lead Contact author. This includes viruses, vaccines, and primer-probe sets. All reagents will be made available on request after completion of a Materials Transfer Agreement.

#### Data and code availability

All data supporting the findings of this study are available within the paper and are available from the corresponding author upon request. This paper does not include original code. Any additional information required to reanalyze the data reported in this paper is available from the lead contact upon request.

## EXPERIMENTAL MODEL AND SUBJECT DETAILS

### Cells and Viruses

Vero cells expressing human ACE2 and TMPRSS2 (Vero-hACE2-hTMPRSS2 [6, 35], gift from Adrian Creanga and Barney Graham, NIH) were cultured at 37°C in Dulbecco’s Modified Eagle medium (DMEM) supplemented with 10% fetal bovine serum (FBS), 10□mM HEPES (pH 7.3), 100□U/mL of Penicillin-Streptomycin, and 10 μg/mL of puromycin. Vero cells expressing TMPRSS2 (Vero-hTMPRSS2) [35] were cultured at 37°C in Dulbecco’s Modified Eagle medium (DMEM) supplemented with 10% fetal bovine serum (FBS), 10□mM HEPES (pH 7.3), 100□U/mL of Penicillin-Streptomycin, and 5 μg/mL of blasticidin.

SARS-CoV-2 (strain 2019-nCoV/USA-WA1/2020) was obtained from the US Centers for Disease Control (CDC) and propagated on Vero-hTMPRSS2 cells. The B.1.621 variant of SARS-CoV-2 (strain hCoV-19/USA/WI-UW-4340/2021) was obtained from a nasal swab isolate and propagated on Vero-hTMPRSS2 cells. Recombinant SARS-CoV-2 with a N501Y and D614G mutations in the S protein of SARS-CoV-2 has been published previously [6] and was propagated on Vero-hTMPRSS2 cells. The virus stocks were subjected to next-generation sequencing, and the S protein sequences were identical to the original isolates. The infectious virus titer was determined by plaque or focus-forming assay on Vero-hACE2-hTMPRSS2 or Vero-hTMPRSS2 cells.

### Pre-clinical vaccine mRNA and lipid nanoparticle production process

A sequence-optimized mRNA encoding prefusion-stabilized Wuhan-Hu-1 (mRNA-1273) SARS-CoV-2 S-2P protein was synthesized *in vitro* using an optimized T7 RNA polymerase-mediated transcription reaction with complete replacement of uridine by N1m-pseudouridine [36]. The reaction included a DNA template containing the immunogen open-reading frame flanked by 5’ untranslated region (UTR) and 3’ UTR sequences and was terminated by an encoded polyA tail. After transcription, the cap-1 structure was added to the 5’ end using the vaccinia virus capping enzyme (New England Biolabs) and vaccinia virus 2’-O-methyltransferase (New England Biolabs). The mRNA was purified by oligo-dT affinity purification, buffer exchanged by tangential flow filtration into sodium acetate, pH 5.0, sterile filtered, and kept frozen at −20°C until further use.

The mRNA was encapsulated in a lipid nanoparticle through a modified ethanol-drop nanoprecipitation process described previously [37]. Ionizable, structural, helper, and polyethylene glycol lipids were briefly mixed with mRNA in an acetate buffer, pH 5.0, at a ratio of 2.5:1 (lipid:mRNA). The mixture was neutralized with Tris-HCl, pH 7.5, sucrose was added as a cryoprotectant, and the final solution was sterile-filtered. Vials were filled with formulated lipid nanonparticle and stored frozen at −20°C until further use. The vaccine product underwent analytical characterization, which included the determination of particle size and polydispersity, encapsulation, mRNA purity, double-stranded RNA content, osmolality, pH, endotoxin, and bioburden, and the material was deemed acceptable for *in vivo* study.

#### Recombinant proteins

Recombinant S, was expressed as previously described [38]. Briefly, a mammalian cell codon-optimized nucleotide sequence coding for soluble S (GenBank: MN908947.3, amino acids 1-1,213) modified to remove the polybasic cleavage site (RRAR to A), but introducing two stabilizing mutations (K986P and V987P, wild-type numbering) and a C-terminal thrombin cleavage site, T4 foldon trimerization domain, and a 6xHIS tag were cloned into mammalian expression vector pCAGGS [39]. Recombinant S was produced in Expi293F cells (ThermoFisher, Cat #A14527) by transfection with purified DNA using the ExpiFectamine 293 Transfection Kit (ThermoFisher, Cat #A14524). Supernatants from transfected cells were harvested 4 days post-transfection, and recombinant proteins were purified using Ni-NTA agarose (ThermoScientific, Cat #88222), then buffer exchanged into phosphate buffered saline (PBS) and concentrated using Amicon Ultracel centrifugal filters (EMD Millipore, UFC903024).

#### Mouse experiments

Animal studies were carried out in accordance with the recommendations in the Guide for the Care and Use of Laboratory Animals of the National Institutes of Health. The protocols were approved by the Institutional Animal Care and Use Committee at the Washington University School of Medicine (assurance number A3381–01). Seven-to-nine week old male 129S2 (strain: 129S2/SvPasCrl, Cat # 287) or female K18-hACE2 transgenic mice (strain: 2B6.Cg-Tg(K18-ACE2)2Prlmn/J, Cat # 34860) were obtained from Charles River Laboratories and Jackson Laboratories, respectively and housed at Washington University. Animals were housed in groups and fed standard chow diet.

Some of the animals were vaccinated via intramuscular (IM) route with 10^8^, 10^9^, or 10^10^ viral particles of fresh or freeze-thawed Ad26.COV2.S in 100 μL of phosphate buffered saline (PBS). The freeze-thawed vaccine was stored at -80°C prior to thawing at room temperature. No difference in serum antibody responses were detected between the fresh and freeze-thawed Ad26.COV2.S vaccine (**Fig S1A-B**). Control animals for the adenoviral vaccine received PBS alone. Twenty-one days and 115 days later, serum samples were obtained, and used for ELISA and virus neutralization assays. Separately, 129S2 mice and K18-hACE2 mice were immunized and boosted with 0.25 or 5 μg of mRNA-1273 or a control mRNA (mRNA-control) vaccine at three week intervals. Twenty-one days after the second immunization, serum was obtained and used for virus neutralization assays.

Following transfer to the enhanced Biosafety level 3 laboratory, the animals were challenged via intranasal route with 10^3^ or 10^5^ PFU of the SARS-CoV-2 N501Y/D614G or B.1.621 variant. Animal weights were measured daily for the duration of the experiment. At different time points after challenge, the animals were sacrificed, and their lungs were collected for virological and histological analysis. The left lobe was homogenized in 1.0 mL of Dulbecco’s Modified Eagle Medium (DMEM), clarified by centrifugation (1,000 x g for 5 min) and used for viral titer analysis by quantitative RT-PCR (RT-qPCR) using primers and probes targeting the *N* gene, and by plaque assay. A nasal wash also was collected, by inoculating 1.0 mL of PBS with 0.1% bovine serum albumin into one nostril and collecting the wash from the other nostril (**Fig 1** and **4**). Alternatively, 0.5 mL of PBS with 0.1% bovine serum albumin was flushed through the nasal cavity after dissecting off the lower jaw (**Fig 2-3**). The nasal wash was clarified by centrifugation (2,000 x g for 10 min) and used for viral titer analysis by RT-qPCR using primers and probes targeting the *N* gene, and by plaque assay.

#### Hamster experiments

Animal studies were carried out in accordance with the recommendations in the Guide for the Care and Use of Laboratory Animals of the National Institutes of Health. The protocols were approved by the Institutional Animal Care and Use Committee at the Washington University School of Medicine (assurance number A3381–01). Five-week old male hamsters were obtained from Charles River Laboratories and housed at Washington University. Five days after arrival, the animals were immunized via intramuscular injection with 10^8^ of 10^10^ viral particles of freeze-thawed Ad26.COV2.S in 100 μL of PBS. Control animals received PBS alone. Twenty-one days later, serum samples were obtained, and the animals were transferred to the enhanced Biosafety level 3 laboratory. One day later, the animals were challenged via intranasal route with 10^3^ PFU of WA1/2020 or B.1.621 variant. Animal weights were measured daily for the duration of the experiment. Four days after challenge, the animals were sacrificed, and their lungs were collected for virological and histological analysis. The left lobe was homogenized in 1.0 mL of DMEM, clarified by centrifugation (1,000 x g for 5 min) and used for viral titer analysis by quantitative RT-PCR using primers and probes targeting the *N* gene, and by plaque assay. A nasal wash was also collected, by inoculating 1.0 mL of PBS with 0.1% bovine serum albumin into one nostril and collecting the wash from the other nostril. The nasal wash was clarified by centrifugation (2,000 x g for 10 min) and used for viral titer analysis by quantitative RT-PCR using primers and probes targeting the *N* gene, and by plaque assay.

## METHOD DETAILS

### Focus reduction neutralization titer assay (FRNT)

Serial dilutions of serum samples were incubated with 10^2^ focus-forming units (FFU) of different strains of SARS-CoV-2 for 1 h at 37°C. Antibody-virus complexes were added to Vero-hTMPRSS2 cell monolayers in 96-well plates and incubated at 37°C for 1 h. Subsequently, cells were overlaid with 1% (w/v) methylcellulose in Eagle’s Minimal Essential medium (MEM, Thermo Fisher Scientific). Plates were harvested 30 h later by removing overlays and fixed with 4% paraformaldehyde (PFA) in PBS for 20 min at room temperature. Plates were washed and sequentially incubated with an oligoclonal pool of SARS2-2, SARS2-11, SARS2-16, SARS2-31, SARS2-38, SARS2-57, and SARS2-71 [40] anti-S protein antibodies and HRP-conjugated goat anti-mouse IgG (Sigma Cat # A8924) in PBS supplemented with 0.1% saponin and 0.1% bovine serum albumin. SARS-CoV-2-infected cell foci were visualized using TrueBlue peroxidase substrate (KPL) and quantitated on an ImmunoSpot microanalyzer (Cellular Technologies).

### Virus titration assays

Plaque assays were performed on Vero-hACE2-hTRMPSS2 cells in 24-well plates. Lung tissue homogenates or nasal washes were diluted serially by 10-fold, starting at 1:10, in cell infection medium (DMEM + 2% FBS + 100□U/mL of penicillin-streptomycin). Two hundred and fifty microliters of the diluted virus were added to a single well per dilution per sample. After 1 h at 37°C, the inoculum was aspirated, the cells were washed with PBS, and a 1% methylcellulose overlay in MEM supplemented with 2% FBS was added. Seventy-two hours after virus inoculation, the cells were fixed with 4% formalin, and the monolayer was stained with crystal violet (0.5% w/v in 25% methanol in water) for 1 h at 20°C. The number of plaques were counted and used to calculate the plaque forming units/mL (PFU/mL).

To quantify viral load in lung tissue homogenates and nasal washes, RNA was extracted from 100 μL samples using E.Z.N.A.^®^ Total RNA Kit I (Omega) and eluted with 50 μL of water. Four microliters RNA was used for real-time RT-qPCR to detect and quantify *N* gene of SARS-CoV-2 using TaqMan™ RNA-to-CT 1-Step Kit (Thermo Fisher Scientific) as described [41] using the following primers and probes: Forward: GACCCCAAAATCAGCGAAAT; Reverse: TCTGGTTACTGCCAGTTGAATCTG; Probe: ACCCCGCATTACGTTTGGTGGACC; 5’Dye/3’Quencher: 6-FAM/ZEN/IBFQ. Viral RNA was expressed as *N* gene copy numbers per mg for lung tissue homogenates or mL for nasal swabs and nasal washes, based on a standard included in the assay, which was created via *in vitro* transcription of a synthetic DNA molecule containing the target region of the *N* gene.

#### Histology

The lungs from SARS-CoV-2 infected and control mice and hamsters were fixed in 10% formalin for seven days. Lungs were embedded in paraffin and sectioned before hematoxylin and eosin staining. Lung slides were scanned using the Hamamatsu NanoZoomer slide scanning system and head sections were imaged using the Zeiss AxioImager Z2 system.

## ELISA

Ninety-six-well microtiter plates (Nunc MaxiSorp; ThermoFisher Scientific) were coated with 100 μL of recombinant SARS-CoV-2 S protein (Wuhan strain) at a concentration of 1 μg/mL in PBS (Gibco) at 4 °C overnight; negative control wells were coated with 1 μg/mL of BSA (Sigma). Plates were blocked for 1.5 h at room temperature with 280 μL of blocking solution (PBS supplemented with 0.05% Tween-20 (Sigma) and 10% FBS (Corning)). Serum from mice and hamsters were diluted serially in blocking solution, starting at 1:100 dilution and incubated for 1.5 h at room temperature. The plates were washed three times with T-PBS (1X PBS supplemented with 0.05% Tween-20), and 100 μL of goat anti-mouse IgG (Southern Biotech Cat #1030-05) diluted 1:2,000 in blocking solution or 100 μL of HRP-conjugated anti-hamster IgG(H+L) antibody (Southern Biotech Cat. #6061-05) diluted 1:500 in blocking solution, was added to all wells and incubated for 1 h at room temperature. Plates were washed 3 times with T-PBS and 3 times with 1X PBS, and 100 μL of 1-step Ultra TMB-ELISA substrate solution (Thermo Fisher Scientific) was added to all wells. The reaction was stopped after 5 min using 100 μL of 1M HCl, and the plates were analyzed at a wavelength of 490 nm using a microtiter plate reader (BioTek).

## QUANTIFICATION AND STATISTICAL ANALYSES

Statistical significance was assigned when *P* values were < 0.05 using GraphPad Prism version 9.3. Tests, number of animals, median values, and statistical comparison groups are indicated in the Figure legends. Analysis of weight change was determined by two-way ANOVA. Changes in infectious virus titer, viral RNA levels, or serum antibody responses were compared to unvaccinated or mRNA-control immunized animals and were analyzed by one-way ANOVA with a multiple comparisons correction, unpaired t-test, or Mann-Whitney test, dependent on the number of comparison and the distribution of the data.

## SUPPLEMENTARY FIGURE LEGENDS

**Figure S1. Effect of an additional freeze-thaw on the immunogenicity of the Ad26.COV2.S vaccine, Related to Fig 1.** (**A**) Neutralizing titer (IC_50_) against WA1/2020 N501Y/D614G of serum obtained from 129S2 mice immunized once with 10^8^ (red symbols), 10^9^ (purple symbols), or 10^10^ (blue symbols) of fresh (solid symbols) or freeze-thawed (open symbols) Ad26.COV2.S. (ns = not significant by unpaired t-test). (**B**) Serum anti-S protein antibody response (EC_50_) in control mice (black symbols), and mice immunized with 10^8^ (red symbols), 10^9^ (purple symbols), or 10^10^ (blue symbols) of fresh or freeze-thawed Ad26.COV2.S (ns = not significant by unpaired t-test). (**C**) Serum anti-S protein specific antibody response (EC_50_) in mice 21 and 115 days after immunization with 10^8^ (red symbols), 10^9^ (purple symbols), or 10^10^ (blue symbols) of fresh or freeze-thawed Ad26.COV2.S. Each symbol represents an individual animal.

**Figure S2. Serum neutralization titer of the B.1.621 variant by sera from immunized mice and Syrian hamsters against, Related to Fig 1-4.** (**A**) Pairwise comparison of the neutralizing titer (IC_50_) against WA1/2020 N501Y/D614G and B.1.621 for individual sera obtained from 129S2 mice immunized once with 10^8^ (red symbols), 10^9^ (purple symbols), or 10^10^ (blue symbols) of fresh (solid symbols) or freeze-thawed (open symbols) Ad26.COV2.S. (ns = not significant by paired t-test). (**B**) Pairwise comparison of the neutralizing titer (IC_50_) against WA1/2020 N501Y/D614G and B.1.621 for individual sera obtained from K18-hACE2 mice immunized twice with 0.25 μg (open symbols) or 5 μg (closed symbols) of mRNA1273 vaccine. (** *P* < 0.01, * *P* < 0.05 by paired t-test). (**C**) Pairwise comparison of the neutralizing titer (IC_50_) against WA1/2020 N501Y/D614G and B.1.621 for individual sera obtained from 129/S2 mice immunized twice with 0.25 μg (open symbols) or 5 μg (closed symbols) of mRNA1273 vaccine. (ns = not significant by paired t-test). (**D**) Pairwise comparison of the neutralizing titer (IC_50_) against WA1/2020 and B.1.621 for individual sera obtained from Syrian hamsters immunized once with 10^8^ (red symbols) or 10^10^ (blue symbols) of fresh (solid symbols) or freeze-thawed (open symbols) Ad26.COV2.S. (**** *P* < 0.0001, *** *P* < 0.001, by paired t-test). Each symbols is an individual animal.

**Figure S3. Histological analysis of lung tissue sections from mRNA-1273 and mRNA-control immunized and K18-hACE2 mice challenged with WA1/2020 N501Y/D614G or B.1.621, Related to Fig 2.** Representative images of 50x, 200x and 400x magnification of hematoxylin and eosin staining of lung sections from K18-hACE2 mice immunized with 0.25 μg (**A**) and 5 μg (**B**) of mRNA-1273 or an mRNA-control (mRNA-CTRL) vaccine and challenged 64 days later with WA1/2020 N501Y/D614G or B.1.621. (**C**) A mock infection is included as a control. Lungs were collected 7 days post challenge, fixed in 10% formalin and paraffin embedded prior to sectioning and staining. The scale bar is 1 mm, 0.25mm and 0.1mm for 50x, 200x and 400x respectively. Representative images are shown from n = 2 per group.

**Figure S4. Histological analysis of lung tissue sections from mRNA-1273 and mRNA-control immunized 129S2 mice challenged with WA1/2020 N501Y/D614G or B.1.621, Related to Fig 3.** Representative images at 50x, 200x and 400x magnification of hematoxylin and eosin staining of lung sections from 129S2 mice immunized with 0.25 μg (**A**) and 5 μg (**B**) of mRNA-1273 or an mRNA-control (mRNA-CTRL) vaccine and challenged 62 days later with WA1/2020 N501Y/D614G or B.1.621 virus. (**C**) A mock infection is included as a control. Lungs were collected 4 days post challenge, fixed in 10% formalin and paraffin embedded prior to sectioning and staining. The scale bar is 1 mm, 0.25mm and 0.1mm for 50x, 200x and 400x respectively. Representative images are shown from n = 2 per group.

