## Supplementary figures and images for "mRNA-1273 and Ad26.COV2.S vaccines protect against the B.1.621 variant of SARS-CoV-2"

### Supplemental Figure 1

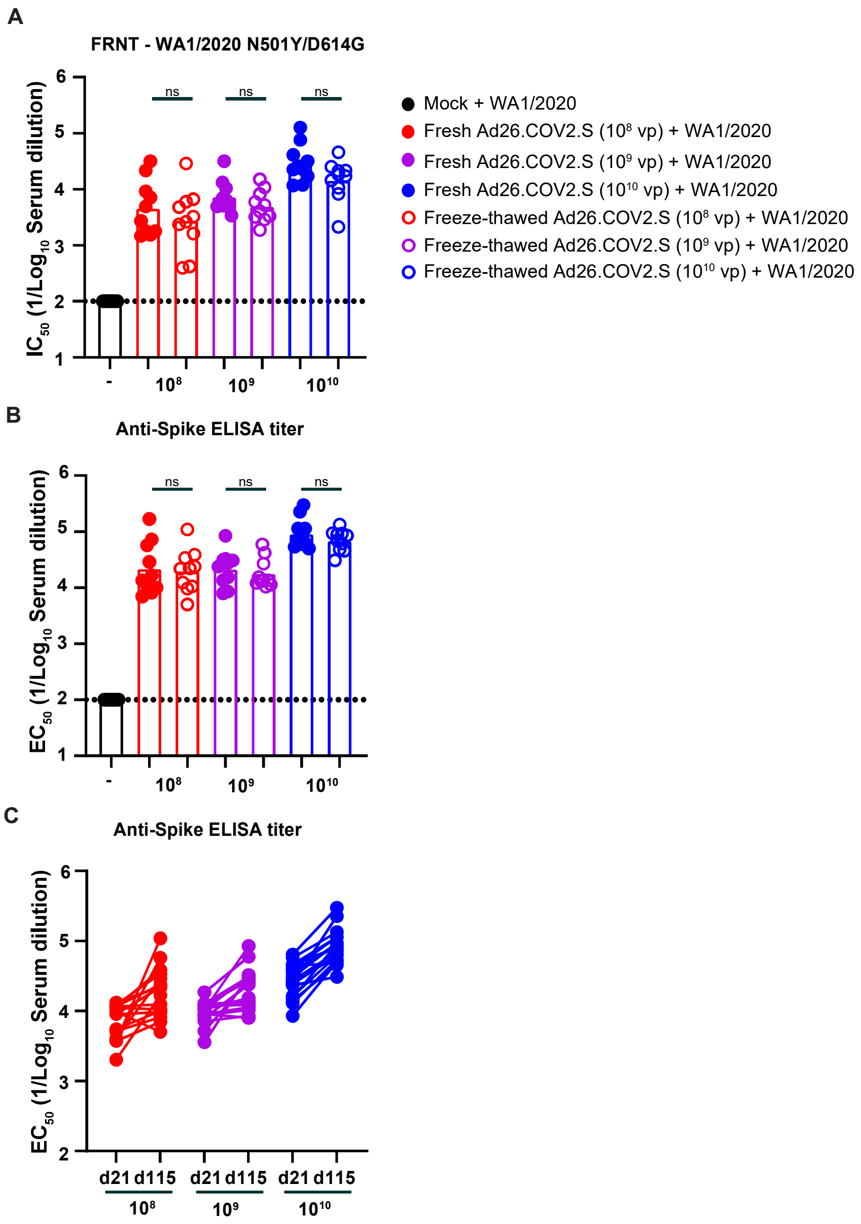

### Supplemental Figure 2

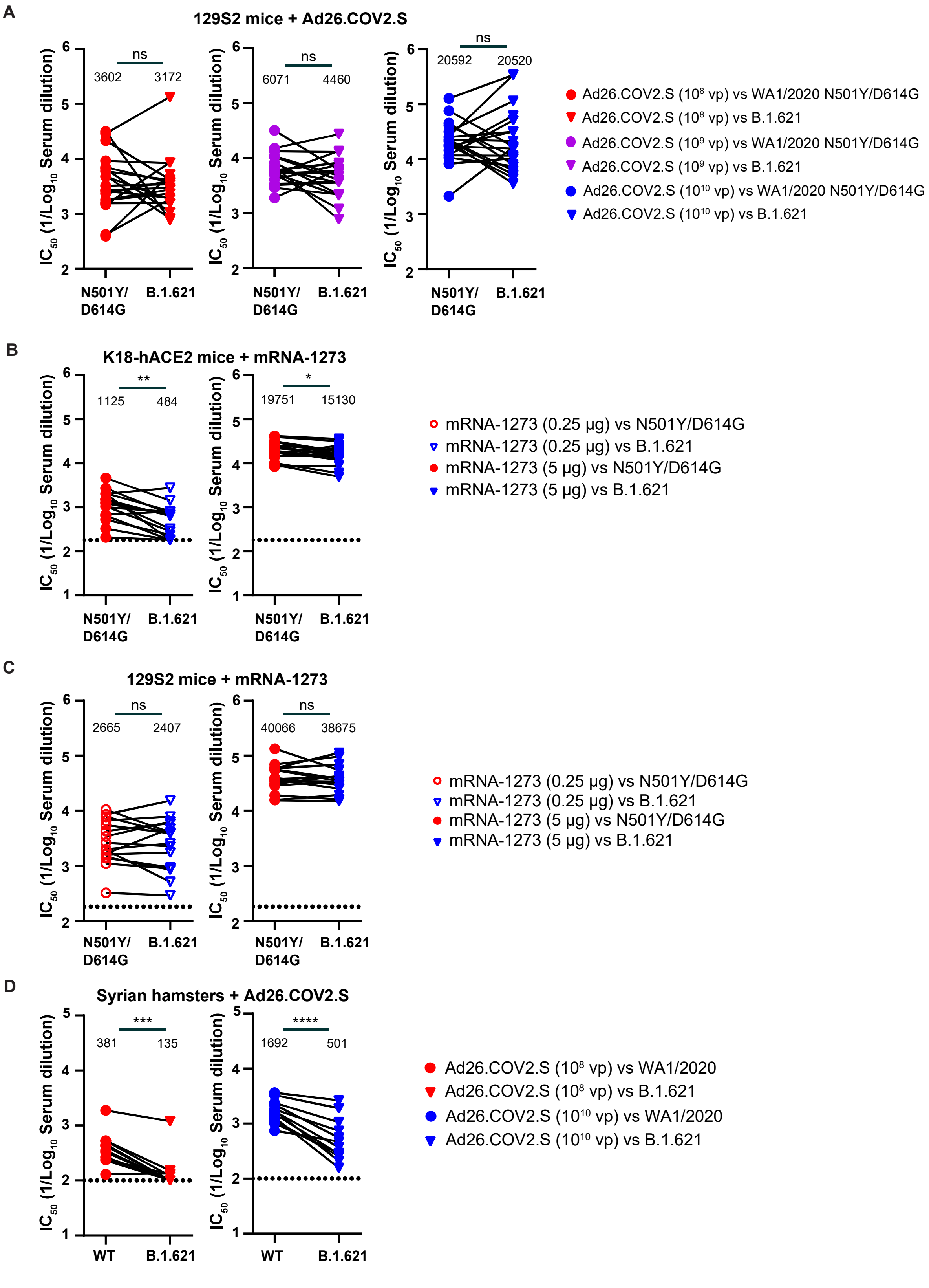

### Supplemental Figure 3

## A. 0.25 μg

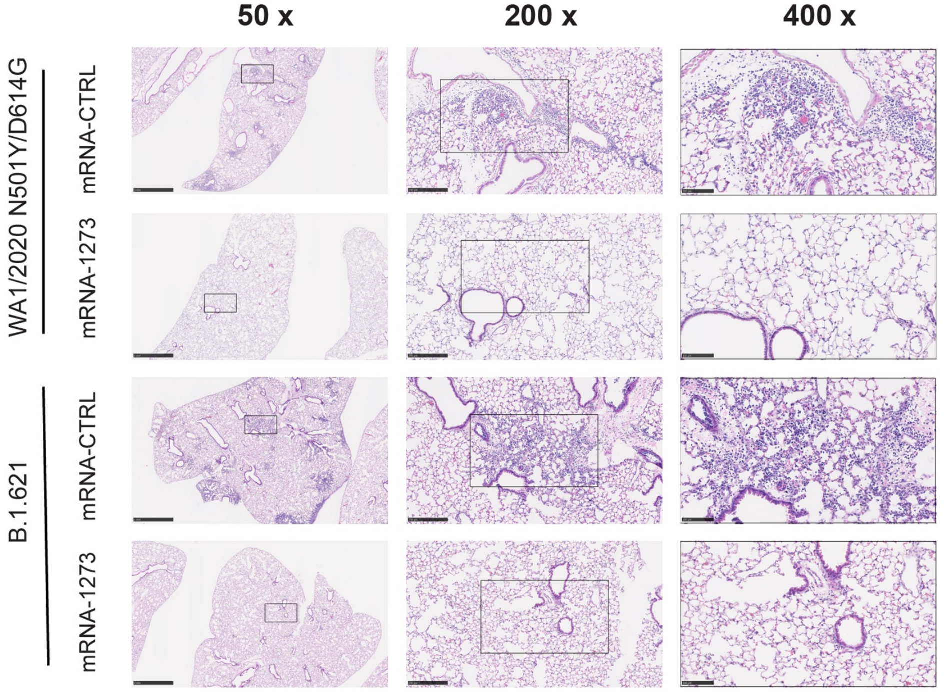


## B. 5 μg

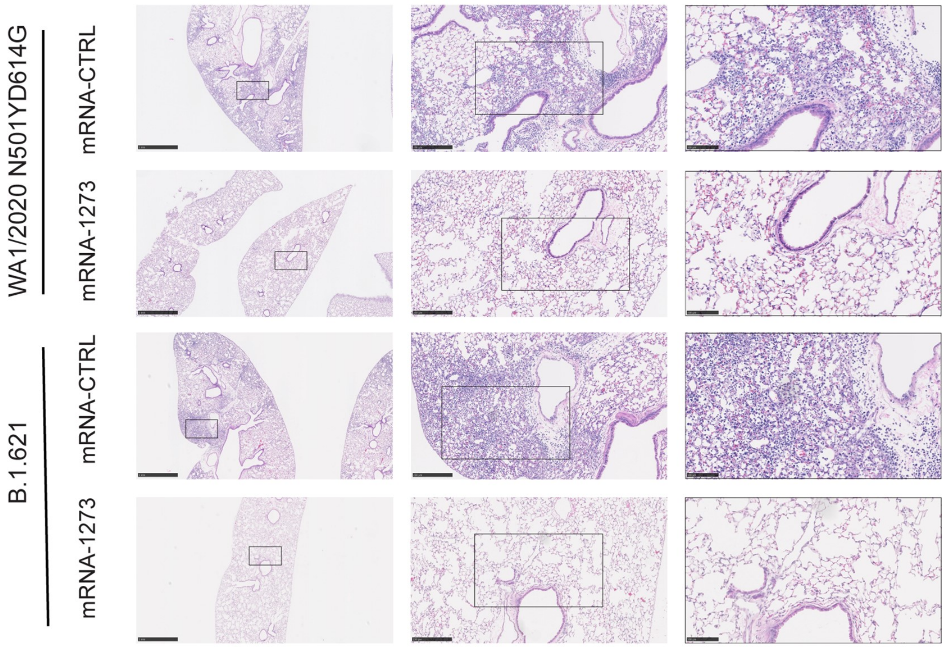


#### C. Mock

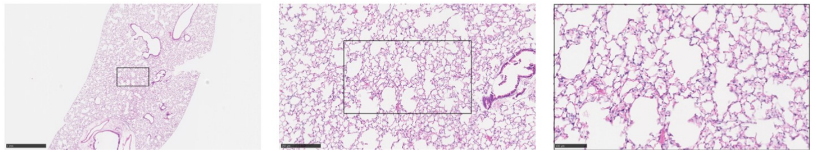

### Supplemental Figure 4

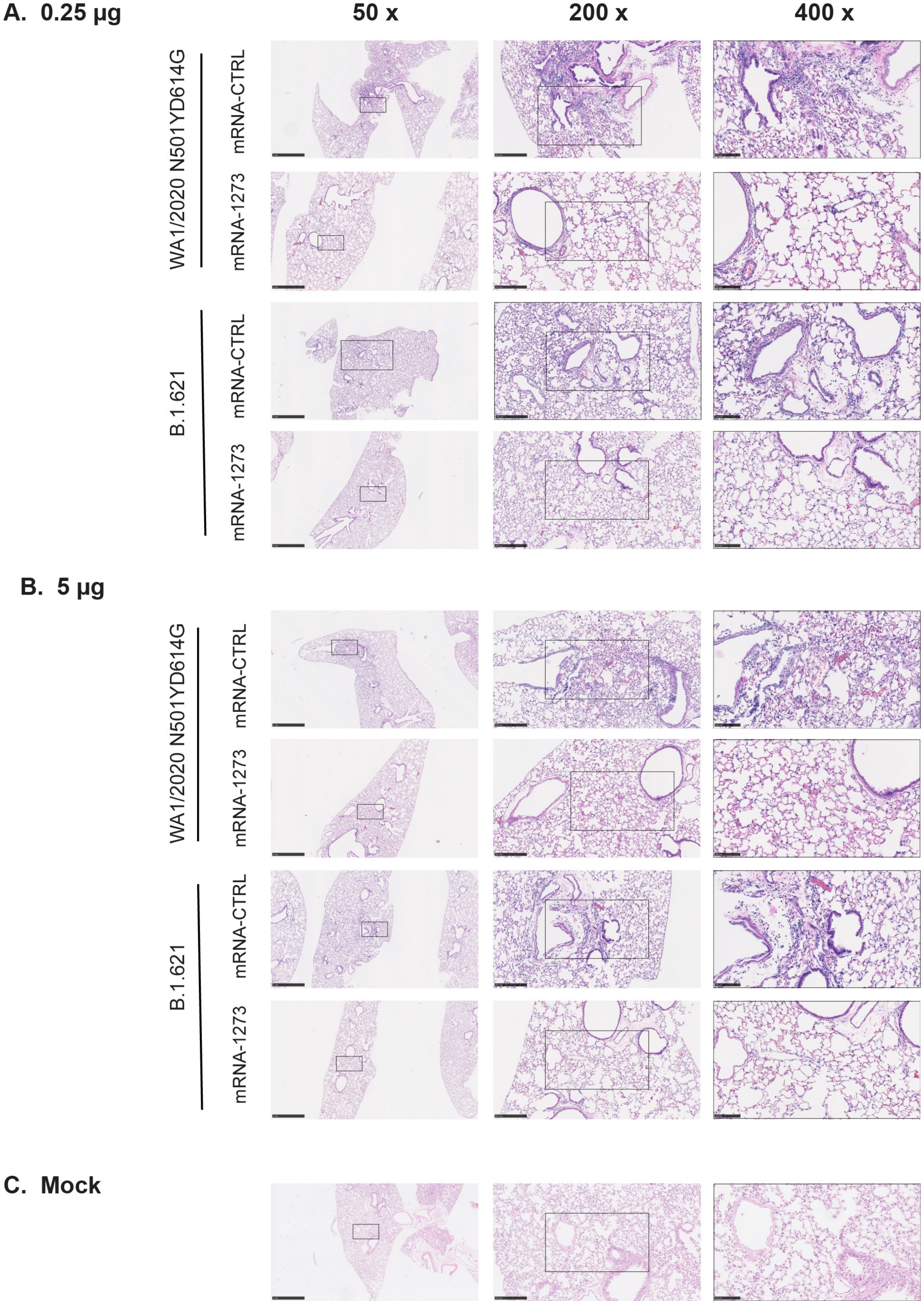
